# DNA isolation protocol effects on nuclear DNA analysis by microarrays, droplet digital PCR, and whole genome sequencing, and on mitochondrial DNA copy number estimation

**DOI:** 10.1101/151126

**Authors:** Elizabeth Nacheva, Katya Mokretar, Aynur Soenmez, Alan M Pittman, Colin Grace, Roberto Valli, Ayesha Ejaz, Selina Vattathil, Emanuela Maserati, Henry Houlden, Jan-Willem Taanman, Anthony H Schapira, Christos Proukakis

## Abstract

Potential bias introduced during DNA isolation is inadequately explored, although it could have significant impact on downstream analysis. To investigate this in human brain, we isolated DNA from cerebellum and frontal cortex using spin columns under different conditions, and salting-out. We first analysed DNA using array CGH, which revealed a striking wave pattern suggesting primarily GC-rich cerebellar losses, even against matched frontal cortex DNA, with a similar pattern on a SNP array. The aCGH changes varied with the isolation protocol. Droplet digital PCR of two genes also showed protocol-dependent losses. Whole genome sequencing showed GC-dependent variation in coverage with spin column isolation from cerebellum. We also extracted and sequenced DNA from substantia nigra using salting-out and phenol / chloroform. The mtDNA copy number, assessed by reads mapping to the mitochondrial genome, was higher in substantia nigra when using phenol / chloroform. We thus provide evidence for significant method-dependent bias in DNA isolation from human brain, as reported in rat tissues. This may contribute to array “waves”, and could affect copy number determination, particularly if mosaicism is being sought, and sequencing coverage. Variations in isolation protocol may also affect apparent mtDNA abundance.

## Introduction

Isolation of DNA is possible in several ways, but often little attention is paid to the protocol, which is not always even reported in detail, with the assumption that the resulting DNA will be a balanced representation of the original source. Any bias in its composition could lead to significant downstream effects on copy number estimation, particularly if mosaicism is being sought, and differential sequencing coverage. Array-based methods have been used to investigate copy number (CN) mosaicism although array “waves” are a recognized problem [1–6], and not fully eliminated bioinformatically [7–10]. Whole genome sequencing (WGS) relative depth of coverage, now frequently used for CN estimation [11], also varies in a wave-like pattern [12–14], which is not fully corrected by PCR-free library construction [15]. Droplet digital PCR (ddPCR) [16] can detect targeted sub-integer changes expected in mosaicism [17] [18]. Bias in DNA isolation has been reported in rat tissues, although CNV mosaicism was first considered as an explanation of the results [12]. To investigate whether DNA isolation bias also occurs in human brain, we analysed DNA isolated with different protocols (with and without spin columns) using the above methods. We found a significant effect of the protocol on downstream results. Care should be given to the selection of DNA isolation method in all applications, with spin columns requiring particular attention. Furthermore, mtDNA copy number determination is influenced by the DNA isolation method chosen [19,20]. We have confirmed this in human substantia nigra, with phenol / chloroform leading to a higher apparent number. Comparison of mtDNA copy number would be prone to error unless the exact same conditions were used.

## Materials and Methods

### DNA samples and isolation

Fresh frozen brain material was provided by the Parkinson’s UK Tissue bank. Donors had given informed written consent. Study of brains from the research tissue bank is approved by the UK National Research Ethics Service (07/MRE09/72). Over the course of this study, we analysed brain DNA from a total of 11 individuals. This included six with Parkinson’s disease (PD), one with incidental Lewy body disease (ILBD; PD-like changes found in autopsy in someone who had not been affected by PD clinically), and four controls. The mean age at death was 79.7 (SD 11.7). Details are provided in table 1. As not all were used for the same experiments, and some were used repeatedly, a summary of the isolation method(s) and experiments performed on each is provided in S1 table.

**Table 1.** Demographic details of individuals whose brains were used.

| Sample ID | gender | age at death | disease duration<br>(years) | Post mortem interval<br>(hours) |
| --- | --- | --- | --- | --- |
| PD1 | m | 63 | 9 | 21 |
| PD2 | m | 69 | 4 | 9 |
| PD3 | m | 73 | 6 | 5 |
| PD4 | m | 68 | 7 | 17 |
| PD5 | m | 78 | 10 | 11 |
| PD6 | f | 83 | 30 | 14 |
| ILBD | f | 104 | - | 10 |
| C1 | f | 78 | - | 23 |
| C2 | m | 82 | - | 48 |
| C3 | m | 90 | - | 12 |
| C4 | f | 89 | - | 13 |

DNA isolation protocols used were the following, following manufacturer instructions unless stated.

1. DNeasy^®^ Blood & Tissue spin column (Qiagen), henceforth referred to as SC. We used approximately 25 mg tissue unless otherwise specified. Brain tissue was cut on dry ice, minced and transferred to a 1.5 ml tube. Buffer ATL (180 μl) was added and the samples were homogenized for 1 min with the IKA Eurostar homogenizer. 20 μl of Proteinase K was added to each sample, and digestion was performed at 56 °C, for 2 hours, or overnight where stated. When digestions were performed overnight, RNase A (4 μl, 100 mg/ml) was also added the next day.
2. Gentra^®^ Puregene^®^ (Qiagen). This relies on the “salting-out” method, which developed from early work showing that DNA, which carries a negative charge, can be recovered using salt solutions of increasing ionic strength in anion-exchange chromatography [21]. It has been used as a non-toxic alternative to phenol / chloroform. Comparisons with spin columns on bone marrow had shown it to yield more DNA, but any possible biases were not assessed [22]. We used approximately 50 mg of brain tissue cut on dry ice, minced and transferred into a 15 ml tube with 3 ml Cell Lysis Solution. We performed further steps according to the protocol for 50-100 mg. We included 15 μl Proteinase K overnight incubation at 55°C as recommended for maximal yield, with subsequent treatment with RNase A, the manufacturer-provided protein precipitation solution, and isopropanol, before 70% ethanol wash.
3. Phenol Chloroform. 450 μL STE buffer and 40 μL 20% SDS were added to 25 mg minced brain sample. After 1 hour incubation at 37°C and vortexing, 20μL Proteinase K were added. The sample was mixed by hand and incubated at 60°C for 4 h. After vortexing, another 20μL Proteinase K were added, mixed by hand, and incubated overnight at 37°C with rotation. The next day samples were centrifuged for 30 minutes and supernatant transferred to clean tubes. 400 μL phenol was added and mixed by hand, followed by 10 minutes on ice, and centrifugation for 2 minutes. The top layer was transferred to a fresh tube. An additional 400 μL phenol was added followed by 5 minutes on ice and centrifugation for 2 minutes. The top layer was removed again and 400 μL of chloroform/isoamyl alcohol (24:1) added and mixed by hand. After centrifuging for 2 minutes, the top layer was transferred to a fresh tube and 2 volumes of cold 95% ethanol and inverted. 4% 3M NaAc was added and the tubes inverted again and placed in -20°C overnight. The next day tubes were centrifuged for 30 minutes, the supernatant was discarded, and 500 μL of 70%EtOH was added. After a final 2 minute centrifugation, the supernatant was discarded, and DNA was air dried and resuspended in 50 μL TE.

We note that there were minor differences in the proteinase K treatment between Puregene (following manufacturer guidelines) and Phenol Chloroform, with a slightly higher initial incubation, and addition of more enzyme with rotation at a lower temperature overnight. We did not use RNase with Phenol Chloroform. Control peripheral blood lymphocyte (PBL) DNA samples were provided by the UCL Institute of Neurology Neurogenetics department.

### Microarray work

We designed a custom 8x60k aCGH array using Agilent e-array software, with ~4,400 probes targeting genes relevant to PD, and their surrounding regions (S2 table). Agilent sex-matched human PBL DNA was used as reference unless indicated otherwise (cat. no: male 5190-4370, female 5190-3797). The recommended 500 ng DNA was used in all cases, to avoid any possibility of variable waves due to unequal DNA amount [7], with hybridisation performed according to manufacturer protocol. Analysis was performed using Agilent Genomic Workbench 7.0. Pre-processing included GC correction (2 kb window size) and diploid peak centralization. The recommended ADM2 algorithm was used, with threshold 6 unless otherwise stated, 5 consecutive probes and 10 kb size needed for a call, and “fuzzy zero” (FZ) long range correction on, unless otherwise specified. All data were mapped to hg19. Isochore graphs were produced by Isosegmenter [23].

We also used the Infinium^®^ CytoSNP-850k Beadchip (Illumina), which is designed for enriched coverage of >3,000 dosage-sensitive genes. Hybridisation was performed according to the manufacturer protocol, using 200 ng DNA. Preliminary analysis was done using BlueFuse Multi 4.1, CytoChip module (Illumina). B allele frequency was estimated by HapLOH [24]. Probe IDs, B allele frequencies, Log R ratios, and AB genotype calls were extracted from BlueFuse output, and AB genotypes were converted to plus strand alleles using allele and strand designations provided by Illumina). We phased the samples using SHAPEIT2, with the Thousand Genomes Project (1KG) haplotypes as a phased reference panel. Specifically, we used the 1KG Phase 1 haplotypes with singleton sites excluded (files downloaded from IMPUTE2 website). Each sample was phased independently using 1KG haplotypes only (SHAPEIT2 option no-mcmc). We applied the hapLOH profiling hidden Markov model using the following parameters: number of event states=1, mean event length=20Mb, event prevalence=0.001, max iterations=100, hapLOH posterior probability of imbalance threshold= 0.5.

### Droplet digital PCR

We performed this on the Bio-Rad QX200 system in 20 μL reactions using 40 ng DNA, ddPCR Supermix, and Biorad-designed commercially available primers (*SNCA*-dHsaCP1000476, *EIF2C1*- dHsaCP1000484, *TSC2*- dHsaCP1000061, *RPP30*- dHsaCP1000485). All were FAM-labelled, except for *RPP30* which was labelled with HEX and used as reference. Restriction digestion using HaeIII (NEB) was performed in tandem with the PCR reaction, by including 2u enzyme in a total of 1μl volume made up with CutSmart buffer. Where specified, DNA was digested in advance (200 ng with 5u enzyme in 10 μl volume), and 1/5 of this was used per ddPCR reaction. Reactions were performed in duplicate. After droplet generation, PCR was performed in the Bio-Rad C1000 Touch Thermal Cycler (95°C for 5 mins, 39 cycles of 95°C for 30 seconds and 60°C 1 min, ending with 98°C for 10 mins). CN was then assessed using the QX200 Droplet Reader and QuantaSoft software (v.1.4.0.99), combining the two replicates of each reaction. Statistical analysis was performed using GraphPad Prism v6.0g, GraphPad Software, CA, USA. For comparison of CN of DNA isolated with different protocols, we first analysed data for normality by the D’Agostino & Pearson omnibus, but this could not be demonstrated due to the small sample size; we therefore compared results using non-parametric tests.

### Whole genome sequencing (WGS)

We prepared dual indexed, paired-end libraries from 2 μg genomic DNA, using TruSeq DNA PCR Free chemistry (Illumina) according to standard protocols. The libraries were sequenced 2x101 bases, in one lane of a Rapid Run flowcell on a HiSeq 2500 (cerebellar DNA), and a single lane of a HiSeq 3000 (substantia nigra). fastq files were trimmed of Illumina adapters and soft clipped to remove low-quality bases (Q>10). Picard (1.75) tools (FastqToSam) were used to convert the fastq files to unaligned BAM files. Reads were aligned to hg19 using Novoalign (v3.02.002), including base score quality recalibration. The generated .bam files were sorted in co-ordinate order using Picard tools and fed into GATK for local realignment around indels. Genome coverage metrics were generated by CollectGcBiasMetrics in Picard, and coverage using CalculateHsMetrics. To calculate chromosome-specific coverage, the chromosome 18 or 19 sequence was used as bait. To estimate the number of mtDNA molecules, we repeated the above steps using the revised mitochondrial genome reference sequence (NC_012920). We then divided the coverage of mtDNA by the coverage of the nuclear genome, and further divided by 2 to correct for the diploid nuclear genome.

## Results and discussion

We initially analysed DNA isolated from cerebellum and frontal cortex (FC) by spin-columns (SC) on aCGH. We noted a consistent wave pattern, more prominent in the cerebellum, even though the cerebellar hybridisations had lower derivative log ratio spread [dLRs] values (S1 fig), and hybridization of the male to female reference DNA used showed no waves (Fig 1A, using chromosome 1 as an example). Several aberrations were called in each sample using the standard threshold of 6 (S1 data; mean 10.7, SD 14.4), of which 1/3 had >10 probes underlying them. Raising the threshold progressively eliminated these; there were 5.6 at threshold 7 (SD 8.0, data S2), 2.7 at threshold 8 (SD 2.8; data S3), 1.7 at threshold 9 (SD 1.6, data S4), and 1.06 at threshold 10 (SD 0.9; data S5). From the 17 calls across all samples at this threshold, 14 were gains at a highly polymorphic 14q32.33 locus. The remaining 3 were a 2 Mb deletion, and two apparent gains, partly overlapping with known CNVs (fig S2). We did not seek to verify these gains.

**Fig 1.**
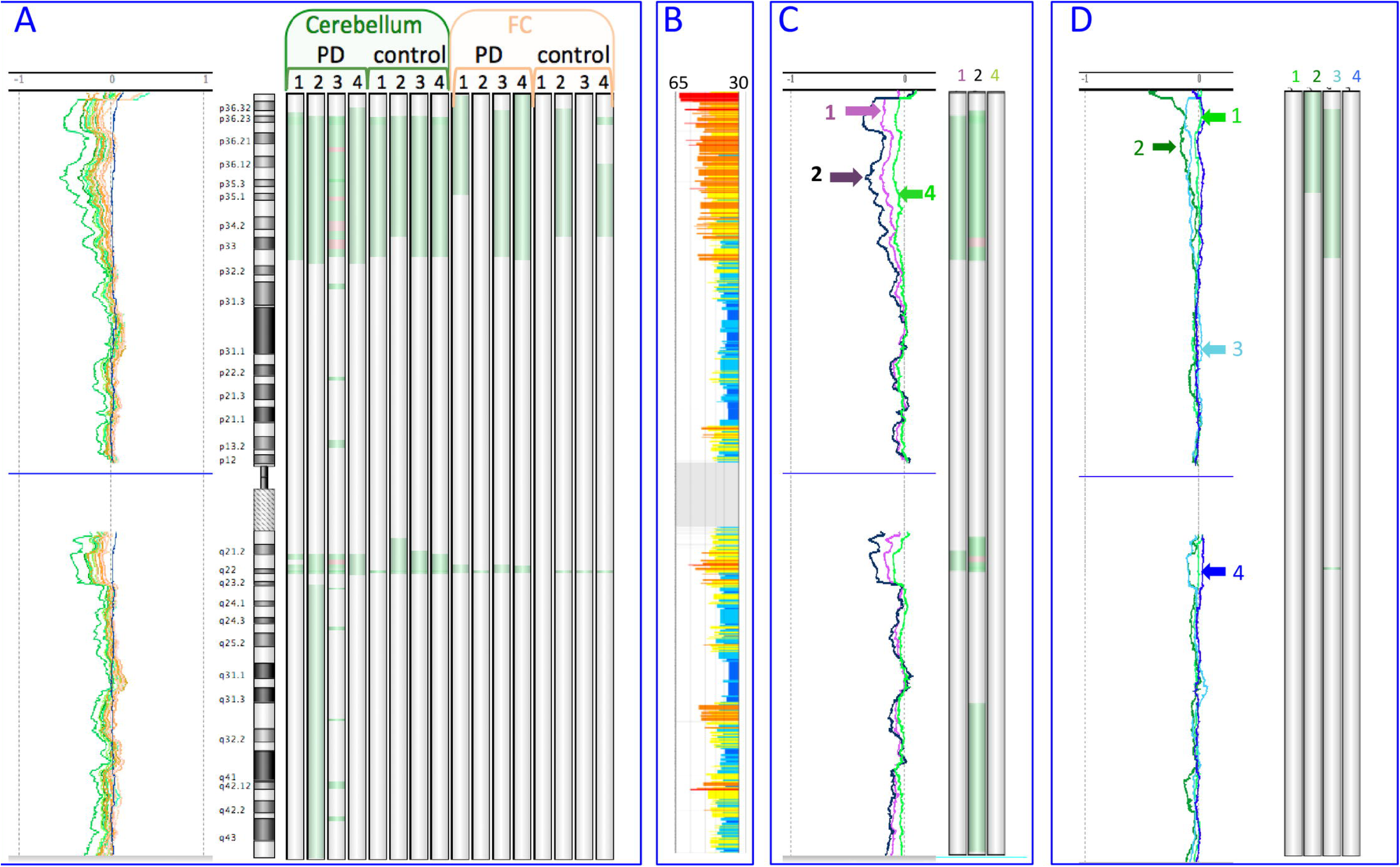
Chromosome 1 in aCGH. The 10 Mb moving average and the aberration calls by ADM2 (after raising threshold to 12, with FZ off) are plotted for each sample. Losses are green, gains are red. (A) Brain DNA hybridised against PBL reference DNA. Cerebellar samples are orange, and FC green. The moving average of a male to female DNA reference hybridisation is also shown (dark blue). (B) Genome isochores. GC content range for each 100 kb isochore is 30-65% (blue to orange). (C) Cerebellar DNA hybridised against FC DNA of the same brain for three PD cases with overnight SC extraction. PD1=purple, PD2=black, PD4=green. Data for PD2 are derived after combining the dye-flip hybridisation pair. (D) Hybridisations between DNA from the same brain as follows. (1-3) Hybridisations of SC-isolated cerebellar DNA, with Puregene-isolated DNA from <u>same</u> cerebellum as reference. (1) PD3, 5 mg SC; (2) PD3, 25mg SC; (3) PD4, 25 mg SC. (4) PD1, Puregene-isolated DNA, cerebellar (test) with FC as reference. Note the absence of waves and losses. This sample combination, but with spin column extraction, had led to waves and losses (PD1 in panel C).

Turning the “fuzzy zero” (FZ) long-range noise correction off, which enhances mosaicism detection [2], and is recommended for this purpose by Agilent in the latest Cytogenomics package, led to more extensive calls at threshold 6, following the “waves”, with apparent losses in GC-rich regions and some gains in GC-poor regions, many of which persisted even after raising the threshold to 12. These often followed the genome GC-content isochores [25] (Fig 1B). There was a clear contrast between chromosome 19, which has the highest gene and CpG island density [26], and displayed negative waves with prominent losses affecting almost its entire length, and the similarly-sized chromosome 18, with the lowest gene density and one of the lowest CpG densities, which showed a mixed picture, with waves in either direction (S2A fig). Chromosome 19 can be problematic on both aCGH [27] and single neuron whole genome amplification [28]. A loss of almost the whole chr19 had indeed been called in one sample by ADM2 with FZ on, but only at threshold 6. To further investigate the apparent excess of subtle losses in cerebellum, we also hybridised cerebellar DNA with FC of the same brain as reference from 3 PD brains, including a dye-flip in one. The wave pattern was still generally present, with several apparent cerebellar relative losses, and reversed by dye flip (fig 1C; S3C–S4 fig).

To investigate the effect of varying the DNA isolation protocol, we isolated cerebellar DNA with SC using overnight proteinase K (rather than 2 hours), starting with approximately 25 or 5 mg tissue in parallel (S3 table), and with the “salting-out” Puregene kit. We noted that the median DNA yield (ng per mg tissue; S3 table) was higher with SC when starting with 5 mg (2201) than with 25 mg (544), and even higher with Puregene (2,784), which was close to the maximum expected (~3,650, based on 6.6 pg DNA per nucleus, and 85 billion cells in a 154 g cerebellum [29]). We then performed aCGH of 25 mg overnight SC isolated DNA for two cerebellar samples, with Puregene-isolated DNA from the same cerebellum as reference; for one of these, we also hybridised a 5mg SC sample to the Puregene sample (fig 1D; S4 fig). The wave pattern in the 25 mg SC samples (2 and 3 in fig 1D) was similar to the original hybridization against PBL DNA, although less pronounced, with some apparent losses called. Waves could therefore be produced even in what was essentially self-hybridisation, although using only 5mg (sample 1 in fig 1D) minimized it. Hybridising Puregene-isolated DNA from cerebellum against FC of one brain (sample 4 in fig 1D and S5 fig) abolished the waves and losses previously seen in the same pair. Our results suggested a differential bias in cerebellum and FC initially, with apparent GC-dependent losses, abolished by using a low amount of tissue and overnight digestion, or Puregene. Using spin columns therefore could lead to incomplete extraction and introduction of a GC-dependent bias, depending partly on the tissue amount used. We used overnight proteinase digestion with Puregene, which should minimize bias, although we cannot exclude the possibility that using a lower tissue amount, or varying the composition of the solution provided by the manufacturer, could be of further help.

To ensure the problem was not limited to our aCGH design, we also analysed freshly isolated DNA (obtained with the original SC protocol) from four control brains (cerebellum in all, and FC in three) on a commercially available SNP array. The logR closely matched the aCGH dLR moving average, with cerebellar losses often called in similar regions to the aCGH negative waves / possible losses (S6 fig), and losses far more frequent than gains (115 v 3 on average; S4 table). We next analysed SNP data using hapLOH [24], which detects regions with significant B-allele frequency (BAF) deviation, and is valuable in the detection of subtle imbalance expected in mosaicism [30]. We found no allelic imbalance, suggesting that the apparent losses affected both chromosomes equally, unlike what one would expect in mosaicism, or heterozygous CNVs (examples in S7 fig). Based on this, we did not feel that the CytoSNP losses called were correct, and we only attempted to validate one by PCR (S7a fig), which was negative (supplementary note), but we cannot exclude the possibility that some were true.

To determine if the isolation protocol could also affect copy number determination by ddPCR, we selected two genes where aCGH suggested negative results (S8 fig); *EIF2C1*, which is also available by the manufacturer as a HEX-labelled reference assay, and *TSC2*, which is implicated in the neurocutaneous disorder tuberous sclerosis, and was within losses in 4/110 frontal neurons in a human single neuron WGS study [31]. The median CN in the original SC samples was less than 2 for both, and lower in the cerebellum than FC, although normal in PBL samples (S9 fig). We compared the results of different protocols on cerebellar DNA (fig 2). The overnight 25 mg SC isolations had higher median CN for both *EIF2C1* (1.77 v 1.33) and *TSC2* (1.64 v 1.31), and the 5 mg SC and Puregene isolation values were even closer to 2 (1.85 and 1.89 for *EIF2C1*; 1.86 and 1.92 for *TSC2*, respectively). There was a highly significant difference in CN between the three conditions tested for all samples (Friedman test p=0.0017 for *EIF2C1* and 0.0046 for *TSC2*), with a significant pairwise difference between the original and Puregene CN values after Dunn’s multiple comparison correction (p=0.0045 and 0.0141 respectively). Modifying the protocol slightly by using a separate restriction digestion step did not alter ddPCR results (S5 table). To determine if ddPCR results for genes outside the negative “wave” regions were influenced by isolation method, we also determined CN for *SNCA,* a gene of major importance in PD, in two cerebellar samples; they were not altered by the isolation method (S6 table). These data, taken together with array results, indicated genuine, protocol-dependent, specific losses during DNA isolation, independent of downstream experiment type.

**Fig 2.**
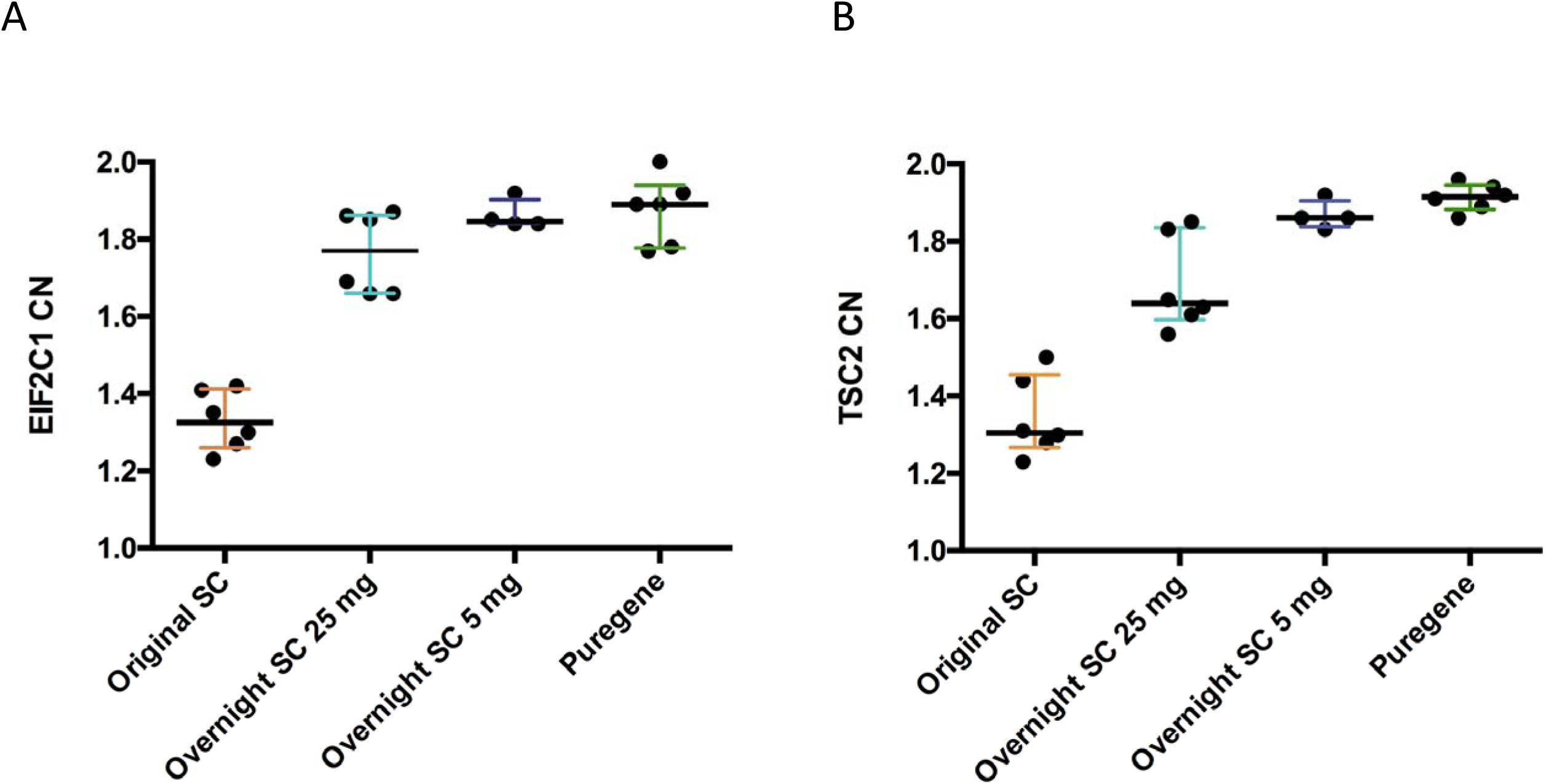
Effect of DNA isolation on copy number determination by ddPCR for cerebellar samples. (A) *EIF2C1* and (B) *TSC2.* The medians and interquartile ranges from the original results, and repeats after overnight SC extractions from 25 mg and 5 mg starting material, and Puregene, are shown. n=6, except 5 mg SC, where n=4.

We next compared low coverage WGS of DNA obtained from the cerebellum with the lowest post-mortem interval (PD3, 5 hours) by a 25 mg SC overnight isolation and by Puregene. We noted a steep decline of coverage with increasing GC content in the SC sample when using 100 kb bins, while the Puregene showed a decline only in the highest GC content (Fig 3). The SC sample showed higher coverage of chr19 compared to chr18, while the Puregene sample had no such bias (ratio 1.4 and 0.97 respectively; S7 table). We then isolated and sequenced DNA from substantia nigra of individuals in parallel using Puregene and the “gold standard” Phenol Chloroform. The Puregene samples revealed a similar GC bias. One of three brains showed the same bias with Phenol Chloroform, while one showed none, even at the highest GC bins (fig S10). The GC “gradient” seen even in the Puregene-isolated samples suggests either that we have not been able to fully remove bias, as in rat tissues [12], or a different GC effect related to the sequencing process, although the Illumina HiSeq provides the most even human genome coverage [15]. The chr18:chr19 coverage ratio did not show major deviation from 1 with either method (S7 table; Phenol 1.03 ± 0.07, Puregene 1.04 ± 0.02), therefore any long-range GC-effect in Puregene and phenol / chloroform may be prominent only in very high GC regions. Phenol / chloroform may have a slight further advantage compared to Puregene, as evidenced by the lack of a 100-kb scale GC gradient on coverage in some cases, although the DNA amount did not allow further experimental comparisons. To determine if WGS GC bias could lead to erroneous copy number calls, even after appropriate corrections, we analysed all data using QDNAseq [32] in 100 kb bins (S11 fig). There were possible losses, but with minimally negative logR, in the SC cerebellar sample, which were absent in Puregene. These would probably be dismissed as noise, although could potentially be misinterpreted as mosaicism.

**Fig 3.**
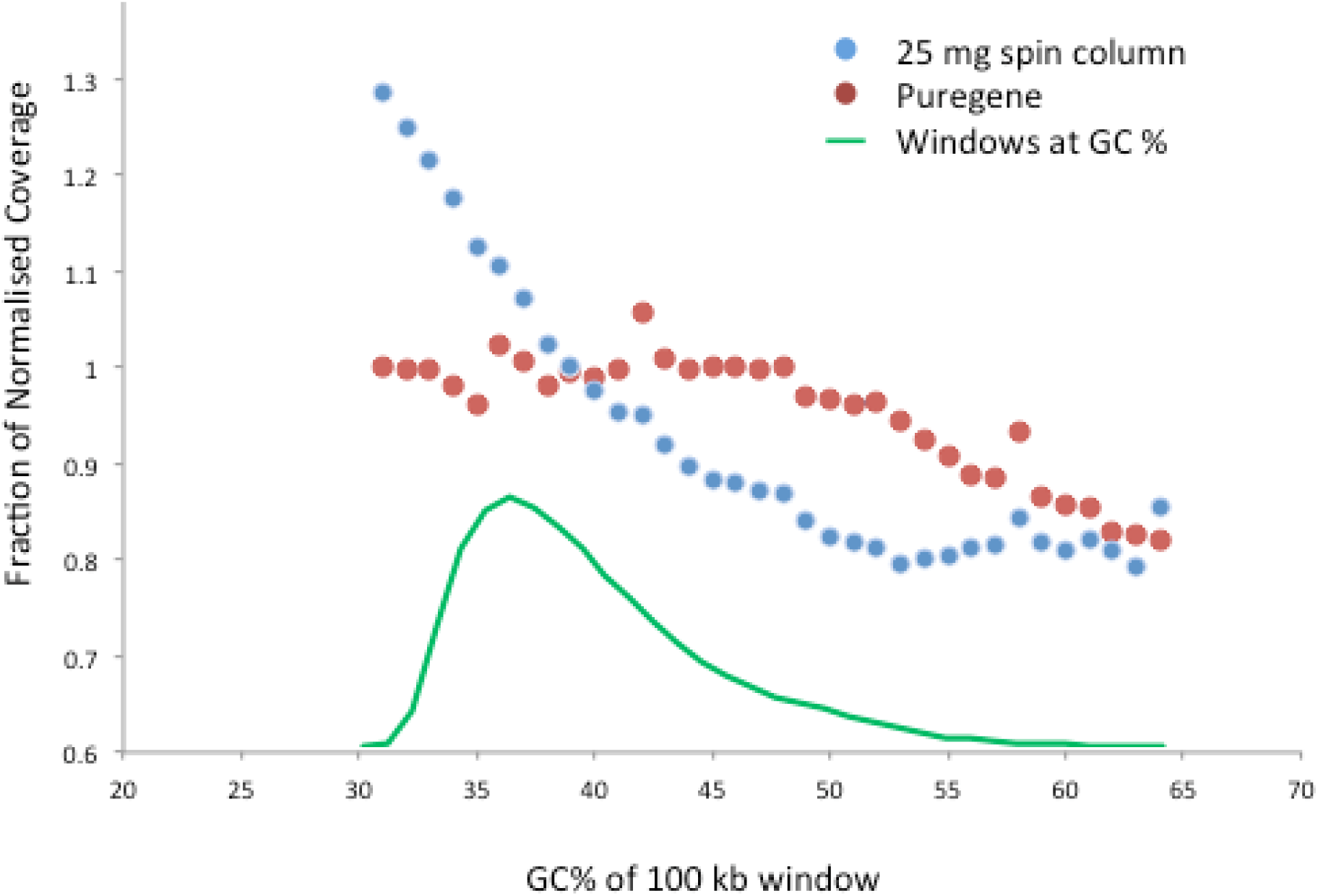
Whole genome sequencing coverage in relation to GC content. The mean normalized coverage per 100 kb window of PD3 cerebellum is shown, after 25 mg overnight SC isolation and Puregene isolation.

We thus demonstrated in human brain that array “waves”, partial losses in ddPCR, and GC-dependent WGS coverage variation, can be modulated, and almost abolished, by variation of the DNA isolation protocol. We have compared the effect of at least two isolation methods on ddPCR for two genes in six cerebellar samples (and on aCGH results in three of them), and on GC-dependent coverage variation in WGS for one of these cerebella, and three substantia nigra samples from different individuals. We therefore believe that we provide strong evidence for uneven GC-dependent DNA extraction, which was recently noted in rat tissues [12], but never before investigated in human tissues to our knowledge, although further studies will help confirm our conclusions. We have not compared PBL DNA isolation, and solid tissues may be most prone to bias. We have data from a single human frozen quadriceps muscle biopsy, from which we isolated DNA with the initial spin column protocol, which we then analysed on the same aCGH design; a similar wave pattern was seen (figure S12). We note a very recent study using only multiple-ligation amplification assay (MLPA) for several fixed human tissues and DNA extraction methods [33]. The number of probes significantly deviating from normality varied between tissues and methods. Although the methodology used was very different to ours, and no information on GC-content of targets was provided, a GC-dependent extraction bias is possible, as acknowledged by the authors.

We found that using longer proteinase K treatment or less material on spin columns, or a non-spin column method, reduced GC-dependent bias. In rats, proteinase K treatment duration had also affected the outcome, but spin columns had not altered results from blood, although this was not examined in other tissues [12]. Strong protein binding to GC-rich DNA regions [12] is a likely mechanism that limits their extraction, particularly if proteinase K digestion is inadequate, or the spin column is saturated. The cerebellum may be more prone to extraction bias may because it is packed with small granule cells, and a greater amount of partly protein-bound DNA in a given tissue mass could result in reduced and more biased overall yield.

Determination of the number of mtDNA copies is of interest in several fields, including PD, where lower mtDNA CN was reported in blood and substantia nigra [34], but with no details on DNA isolation, and cancer, where batch effects were corrected bioinformatically, but remained unexplained [35]. Although traditionally done by qPCR, it is now possible to determine the number of mtDNA molecules in a preparation by the ratio of sequencing reads mapping to the nuclear versus mitochondrial genome [36-37]). We therefore determined this for each sample, from the bulk DNA isolation, without seeking to specifically isolate mtDNA. We then compared the results obtained by different isolation methods (table 2). For the nigra samples, phenol led to a higher number than Puregene (average increase 2.51-fold, SD 0.71). This is consistent with a previous report that organic solvent extraction results in mtDNA enrichment [20]. As we did not use RNase with Phenol, but we did as per the standard protocol with Puregene, we cannot comment on any possible effect of this, although the potential higher mtDNA recovery when omitting RNase may only apply to spin columns [20]. The mtDNA number is similar to a human brain DNA phenol isolation report [19], although much lower than claimed elsewhere [34].

**Table 2.** Effect of DNA isolation on mtDNA copy number estimated by sequencing.

| Source | Isolation method | mtDNA copy number | Ratio |
| --- | --- | --- | --- |
| PD5 SN | Ph:Chl | 1380 | 1.72 |
|  | Puregene | 802 |  |
| PD6 SN | Ph:Chl | 2398 | 2.73 |
|  | Puregene | 877 |  |
| ILBD SN | Ph:Chl | 1225 | 3.08 |
|  | Puregene | 397 |  |
| PD3 CER | SC | 508 | 0.98 |
|  | Puregene | 518 |  |

The ratio of the number estimated for each sample with different isolation methods is shown. Ph:Chl = Phenol / chloroform.

Our results highlight the often overlooked effects of DNA isolation on copy number determination, sequencing coverage variation, and mtDNA copy estimation. Array and sequencing “waves” may be largely due to isolation-induced relative losses. Raising the ADM2 threshold, and keeping the “fuzzy-zero” correction, reduces false positive calls, although may not eliminate them unless high values are used at the expense of sensitivity. Further studies will be helpful for further validation, and detailed assessment in other tissues, but we believe that studies should carefully select and fully report the DNA isolation protocol. For spin columns, the amount of tissue loaded, and the proteinase digestion duration, might require optimisation, and avoiding spin columns may sometimes be preferable. Comparing WGS coverage of chromosomes with different GC content, or performing selective ddPCR, as we have done, can help exclude major GC bias. When comparing different samples, the same protocol should be followed. Suspected CN mosaicism should be confirmed by allelic imbalance, direct visualization by FISH, or breakpoint demonstration. mtDNA number comparisons should be treated with caution unless the exact same conditions were used.

## Acknowledgements

Tissue samples and associated anonymized data were supplied by the Parkinson’s UK Tissue Bank, funded by Parkinson’s UK, a charity registered in England and Wales (258197) and in Scotland (SC037554). We are grateful to Dr Udo Koehler of MGZ Medical Genetics Centre for performing the Beadchip hybridization, the UCL Institute of Neurology sequencing facility, and to all patients and controls who donated their brains to research.

## Supporting information

**S1 fig. Derivative Log Ratio spread (dLRs) values of aCGH brain to PBL reference DNA hybridisations.** CER= cerebellum, FC = frontal cortex. The median and interquartile ranges are shown.

**S2 fig. CNVs called in only one sample with ADM2 threshold 10, FZ on.** See also data S5. The chromosomal location, individual probe DLR with region of call highlighted, and CNV list in this region are shown for each.

**S3 fig. aCGH results for brain DNA of chromosomes 19 (above), and 18 (below).** Analysis by ADM2 (FZ off). The ADM2 threshold is 12 for chr.19, and 6 for chr.18, as most changes were not visible at higher thresholds. 5 Mb moving averages are shown

a. Cerebellum and FC DNA with PBL DNA as reference. Arrows show gains in low GC regions.
b. The human GC content isochore plot (orange=high, blue=low; range 30-65%).
c. PD cerebellar DNA with FC from same brain as reference for PD1, 2, and 4. For PD2, analysis of the combined dye-flip pair is shown. Note that for chr18, in the arrowed low GC region, even the two samples where gains were not called have a slightly positive moving average.

**S4 fig. Dye-flipped hybridisations of PD2 cerebellum and FC DNA.**

(1) Cerebellum (test) v FC (reference), red. (2) FC (test) v cerebellum (reference), dye flip specified during data import, blue. (3) Male to female reference PBL DNA hybridisation, brown, for comparison. Moving averages are shown over 10 Mb for chr.1, and 5 Mb for chr.18 and 19.

**S5 fig. Chromosome 19 (above) and 18 (below) in aCGH analysis of additional DNA extractions with different protocols.** Analysis by ADM2 (FZ off, threshold 12 for chr.19, 6 for chr.18), with 5 Mb moving averages, and GC isochores; range 30-65%). (1-3): Hybridisations of spin column-extracted cerebellar DNA, with Puregene extracted DNA from same cerebellum as reference.

(1) PD3, 5 mg spin column extraction; (2) PD3, 25mg spin column extraction; (3) PD4, 25 mg spin column extraction.

(4) PD1, Puregene DNA, cerebellar, with FC as reference. Note the absence of waves and losses, unlike the same combination but after SC isolations, shown in fig S3C, sample 1).

**S6 fig. Detailed comparison of genome-wide calls by aCGH and SNP array in samples C1 (A) and C2 (B).** These samples had the highest number of losses on SNP array. The five columns for each chromosome are as follows:

1. Deletion calls by SNP array (red dots / bars)
2. Common deletion calls. Yellow lines / bars represent areas called as losses by SNP array and aCGH (using ADM2, threshold 8, FZ off)
3. aCGH (ADM2, threshold 8, FZ off). Losses are green, gains are red.
4. aCGH (ADM2, threshold 12, FZ off).
5. aCGH (ADM2, threshold 12, FZ off). An additional filter was used to filter calls that have a level of <15% gain or loss. Note that almost all the losses at these settings are also called by the SNP array.

The rare gains on SNP array are shown as green dots between columns 2 and 3, and highlighted with a blue arrow.

**S7 fig. Examples of chromosome 1 SNP array losses.** The left hand column shows the relevant data including **with B allele frequency comparison to aCGH** from **C1 cerebellum**. The right hand column shows the SNP logR over this region in selected other three samples where it was also somewhat negative, although losses were not always called.

A. Loss around *FBXO42* (chr1:16,619,350-16,773,880; 154.5 kb). This was examined further by PCR, and not confirmed (see supplementary note).
B. Loss in 1q22 region (chr1: 155,540,660-155,819,657; 279 kb).

**S8 fig. aCGH results around ddPCR target genes in all hybridisations using original brain SC isolations.**

Probe dLRs in each hybridisation are shown, grouped by type, with ddPCR target location indicated by a blue line. For PD2 Cerebellum to FC, the combined the dye-flip data were used.

A. EIF2C1, 325 kb shown (chr1:36199314-3652540)
B. TSC2, 144 kb shown (chr16:2027153-2172009)

**S9 fig. Copy number of EIF2C1 and TSC2 determined by ddPCR in cerebellum (CER), frontal cortex (FC), and peripheral blood leucocytes (PBL).** The median and interquartile ranges are shown in all cases. (a) EIF2C1. CER and FC from four brains, CER only from another three, and three control PBL DNA samples. Kruskal-Wallis p < 0.001 (b) TSC2. CER and FC from three brains, and cerebellum only from another four, and four control PBL DNA samples. Kruskal-Wallis p=0.0001

**S10 fig. WGS coverage in relation to GC content for the three SN samples after Phenol / Chloroform and Puregene isolation.**

The mean normalized coverage per 100 kb window is shown (y-axis) and the % content of each window (x-axis). The base quality for each GC content is also shown.

**S11 fig. QDNAseq analysis of Cerebellar DNA WGS data with different isolation methods in 100 kb bins.** (A) Phenol / Chloroform. (B) Puregene. The total PE read number was 97,978,308 for SC, and 62,120,676 for Puregene. The right-hand figure in A shows the results after downsampling to 62,115,269 reads, done with Picard DownsampleSam (strategy=high accuracy). The estimated minimum standard deviation due purely to read counting (Eσ) and the observed standard deviation (σ_Δ_) are shown. The y-axes show the log ratio (left) and probability assigned to the aberration called (right). The observed losses in A had a minimally negative log ratio, and are indicated by arrows for clarity. The same analysis of the SN WGS data revealed no aberrations (data not shown).

S12 fig. aCGH analysis of a muscle sample isolated by spin column. The moving averages are shown for chr1 (over 10 Mb), and 18 and 19 (5 Mb). Aberrations called with FZ off are highlighted (threshold 12 for chr1 and 19, 6 for chr18).

**S1 table. Summary of all samples and experiments. CER= cerebellum. ON= overnight. S/C= spin column. PG= puregene. Ref= used as reference DNA in aCGH. Individual experiments explained in text.**

**S2 table. aCGH custom design targets and probes.** The fragile site within which *SNCA* is located and its flanking regions were included (chr4:87-97 Mb), as constitutional CNVs may involve most of this.

**S3 table. Effect of DNA extraction protocol on yield and purity of cerebellar DNA.** For each sample, the exact tissue mass used in spin column extractions (in mg), the DNA yield (in ng DNA per mg tissue), and the 260/280 ratio are shown. Yield mean and SD are shown at bottom. ANOVA for yield in four samples where all protocols were used: p=0.039.

**S4 table. Summary of CytoSNP calls in each sample.**

**S5 table. *TSC2* CN in ddPCR performed with and without restriction digestion as a separate step.**

Two control cerebellar samples analysed, with the standard protocol, and with restriction digestion as a separate step (“pre-digested”), in DNA extracted with spin columns overnight, or Puregene.

**S6 table. *SNCA* CN by ddPCR in two control cerebellar samples, using DNA extracted with different protocols.**

**S7 table. WGS summary results of samples isolated with different methods.**

**S1-S5 data.** All the brain DNA aberrations reported against PBL reference DNA with ADM2 at varying thresholds, FZ on. Thresholds are as follows: S1-6, S2-7, S3-8, S4-9,S5-10.

**Supplementary note. Attempted PCR validation of a CytoSNP-called deletion.**

